# Rab35 and glucocorticoids regulate APP and BACE1 trafficking to modulate Aβ production

**DOI:** 10.1101/2021.05.10.443354

**Authors:** Viktoriya Zhuravleva, João Vaz-Silva, Mei Zhu, Patricia Gomes, Joana M. Silva, Nuno Sousa, Ioannis Sotiropoulos, Clarissa L. Waites

## Abstract

Chronic stress and elevated glucocorticoids (GCs), the major stress hormones, are risk factors for Alzheimer’s disease (AD) and promote AD pathomechanisms, including overproduction of toxic amyloid-β (Aβ) peptides and intraneuronal accumulation of hyperphosphorylated Tau protein. The latter is linked to downregulation of the small GTPase Rab35, which mediates Tau degradation via the endolysosomal pathway. Whether Rab35 is also involved in Aβ overproduction remains an open question. Here, we find that hippocampal Rab35 levels are decreased not only by stress/GC but also by aging, another AD risk factor. Moreover, Rab35 negatively regulates Aβ production by sorting amyloid precursor protein (APP) and β-secretase (BACE1) out of the endosomal network, where they interact to produce Aβ. Interestingly, Rab35 coordinates distinct intracellular trafficking events for BACE1 and APP, mediated by its effectors OCRL and ACAP2, respectively. Finally, we show that Rab35 overexpression prevents the amyloidogenic trafficking of APP and BACE1 induced by high GC levels. These studies identify Rab35 as a key regulator of APP processing and suggest that its downregulation may contribute to stress- and AD-related amyloidogenesis.

## Introduction

Alzheimer’s disease (AD) is the most common age-related neurodegenerative disease and cause of dementia. While aging constitutes the highest risk factor for AD ^1^, emerging evidence indicates that chronic stress and high circulating levels of glucocorticoids (GCs), the main stress hormones, are also risk factors for the disease ^2–5^. Consistent with epidemiological studies, recent work has shown that chronic unpredictable stress and GCs induce AD-like pathology in animal models. These include misprocessing of amyloid precursor protein (APP), leading to overproduction of toxic amyloid-beta (Aβ) peptides, as well as the accumulation and hyperphosphorylation of Tau protein, leading to its synaptic mislocalization and downstream synaptic loss and memory impairment ^5–11^.

In previous work, we showed that GC-induced Tau pathology is precipitated in part by transcriptional downregulation of the small GTPase Rab35 ^12^, a master regulator of endosomal protein trafficking. In particular, we found that Rab35 promotes Tau degradation via the endolysosomal pathway and that its downregulation by GCs leads to intraneuronal Tau accumulation ^12^. AAV-mediated overexpression of Rab35 in the rodent hippocampus was sufficient to prevent GC-induced Tau accumulation and downstream dendrite and spine loss ^12^. These findings demonstrate that GCs precipitate Tau pathology by disrupting Rab35-mediated endolysosomal trafficking. In addition to its role in Tau degradation, Rab35 also mediates other protein trafficking events, including retrograde trafficking of mannose-6-phosphate receptors from endosomes to the trans-Golgi network ^13^ and recycling of T-cell receptors and cell adhesion molecules to the plasma membrane (PM) in a pathway that operates in parallel with Rab11-mediated endosomal recycling ^14–17^. Given the multiple trafficking events regulated by Rab35, it is possible that GC-mediated downregulation of Rab35 could also impact APP trafficking and contribute to stress-induced Aβ production.

Trafficking of APP and β-secretase1 (BACE1) into the endosomal network is an essential step in Aβ production ^18,19^. BACE1 mediates the rate-limiting cleavage step in the amyloidogenic processing of APP into Aβ, and endosomes are major sites of Aβ generation due to their optimal acidic pH for BACE1 activity ^19^. Intriguingly, many of the recently identified genetic risk factors for late-onset AD are linked to endosomal protein trafficking, and have been shown to induce endosomal dysfunction and prolong the residence times of APP and/or BACE1 in endosomes ^20,21^. The ~60 member family of Rab GTPases are key regulators of endosomal protein trafficking and have been implicated in the etiology of AD as well as other neurodegenerative diseases ^22–25^. Indeed, a subset of Rabs, including Rab35, were identified as regulators of APP processing and Aβ production in a loss-of-function screen of Rab GTPases performed in non-neuronal cells ^22^. However, it is yet unclear whether Rab35 regulates APP or BACE1 trafficking and Aβ production in neurons, and if so, how this function is impacted by stress/GCs.

Here, we demonstrate that Rab35 levels decrease in the hippocampus in response to chronic stress and aging, both AD risk factors, suggesting that Rab35 reduction may be a precipitating factor for AD-related pathomechanisms. Consistent with these observations, we show that Rab35 inhibits Aβ production in neurons by stimulating APP and BACE1 trafficking out of the endosomal network. Interestingly, our findings indicate that Rab35 regulates distinct trafficking steps for APP and BACE1, mediated by distinct Rab35 effectors (ACAP2 and OCRL, respectively). Finally, we show that high GC levels alter the endosomal trafficking of APP and BACE1 to increase their interaction and that Rab35 overexpression counteracts these GC-induced effects. Together, these findings implicate Rab35 as a negative regulator of amyloidogenic APP processing and suggest that its downregulation could contribute to Aβ production during aging or after prolonged exposure to stressful conditions.

## Results

### Rab35 levels are decreased by aging and under chronic stress

Our previous study showed that treatment with high levels of glucocorticoids (GCs) suppresses Rab35 expression in hippocampal neurons *in vivo* ^12^. To test whether chronic stress has a similar effect, we monitored Rab35 levels in the hippocampi of rats subjected to a chronic unpredictable stress paradigm for 4 weeks. We found that hippocampal Rab35 levels were decreased by ~60% in stressed rats compared to controls (Fig. 1A-B) and that this decrease was not observed for other Rabs associated with endocytic protein trafficking (Fig. 1A-B), in line with our previous findings ^12^. In parallel, we monitored hippocampal Rab levels in animals infused with Aβ, a procedure that is widely used to model early AD neuropathology in rodents and primates ^26–28^. Infused Aβ interact with full-length APP to further stimulate Aβ production ^6,29–31^ and circumvent the need to express human APP containing mutation(s) that impact its localization and trafficking. Importantly, levels of Rab35, but not most other endocytic Rabs, were similarly decreased in Aβ-infused hippocampi (Fig. 1A-B). Since aging is the greatest risk factor for AD and previous studies report increased amyloidogenic APP processing in the aged brain ^32,33^, we also compared Rab35 expression levels in hippocampi of young (4 month-old) versus aged (22-24 month-old) rats. Again, we observed a significant (~25%) decrease in hippocampal Rab35 levels in the aged animals (Fig. 1C-D), indicating downregulation of Rab35 expression in this brain region during aging. To determine whether boosting hippocampal Rab35 levels could inhibit the amyloidogenic processing of APP in aged animals, we injected 17-19 month-old rats with AAV8 to express EGFP or EGFP-Rab35 in the dorsal hippocampus. Intriguingly, we found that Rab35 overexpression decreased the levels of APP C-terminal fragments (CTFs) relative to fulllength APP (Fig. 1E-F). These results suggest that Rab35 inhibits APP amyloidogenic processing and that its reduction during aging and/or stress could contribute to the increased Aβ production observed under these conditions.

**Figure 1.**
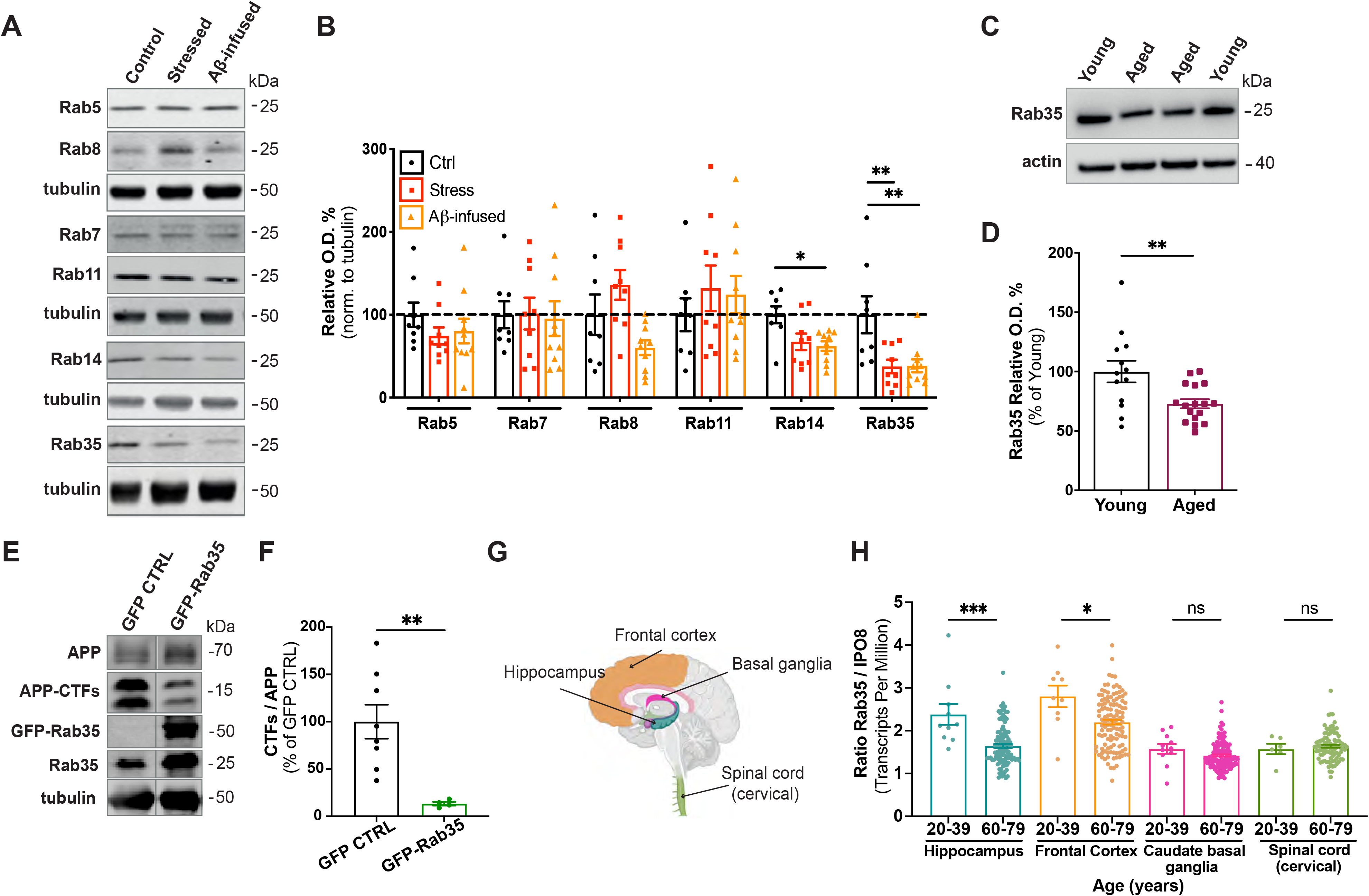
Hippocampal Rab35 levels are decreased by chronic stress and aging. A-B) Representative immunoblots and quantification of Rab protein levels in the hippocampus of control (Ctrl), stressed, and Aβ-infused rats. Blots were probed for the noted Rabs and tubulin, with values normalized to tubulin and expressed as % of the control condition (dotted line). Rab14 and Rab35 are reduced by Aβ, but only Rab35 is significantly reduced by both stress and Aβ (**P_Rab35 CON vs. stressed_ =0.0075, P_Rab14 CON vs. Aβ-infused_ =0.0228, P_Rab35 CON vs. Aβ-infused_ =0.0069; one-way ANOVA, Dunnet post hoc analysis, n=8–10 animals/condition). C-D) Representative immunoblots and quantification of Rab35 levels in the hippocampus of young (4 month-old) and aged (22-24-month-old) rats. Blots were probed for Rab35 and actin, with values normalized to actin and expressed as % of young animals. Rab35 expression is decreased in aged animals (**P=0.0056; unpaired t-test, n=13–17/condition). E-F) Representative immunoblots and quantification of APP C-terminal fragments (CTFs) relative to full-length APP in the dorsal hippocampus of rats bilaterally injected with AAV-GFP or AAV-GFP-Rab35. Blots were probed for APP, Rab35, and tubulin, with values normalized to tubulin and expressed as % of GFP control condition. Rab35 expression significantly decreases the ratio of CTFs to full-length APP (**P=0.0019; Welch’s unpaired two-tailed t-test, n=4-8 animals/condition). G) Schematic diagram of human brain areas used for the analysis of transcriptome data from the Genotype-Tissue Expression (GTEx) project, created with BioRender. H) Quantification of Rab35 mRNA transcripts from individuals ages 20-39 years and 60-79 years, normalized to IPO8 (*P=0.0155, **P=0.0003; Mann-Whitney test, n=6-135 samples/condition), showing the reduction of Rab35 levels in the hippocampus and frontal cortex, but not basal ganglia and spinal cord, of 60-79 year-old individuals. All numeric data represent mean ± SEM.

To investigate whether aging similarly impacts Rab35 levels in the human brain, we analyzed gene expression data from the Genotype-Tissue Expression (GTEx) Portal, a collection of data from non-diseased human tissue (https://gtexportal.org/home/). Transcript levels were normalized to those of IPO8, identified by RefFinder ^34^ as the most stable gene across ages and brain regions of this data set. Intriguingly, we found that Rab35 transcripts were decreased in the hippocampus and frontal cortex from individuals 60-79 years of age compared to those 20-39 years of age (Fig. 1G-H). These changes were not seen in basal ganglia or cervical spinal cord, tissues not implicated in stress- and AD-related pathology (Fig. 1G-H). Together, these findings suggest that Rab35 expression also decreases during human aging in the hippocampus and frontal cortex, brain regions impacted in AD and by chronic stress ^1,35^.

### Rab35 is a negative regulator of APP-BACE1 interaction and Aβ production

A previous study identified Rab35 as a negative regulator of Aβ production in a loss-of-function screen of Rab GTPases in non-neuronal cells ^22^. To better understand whether and how Rab35 regulates Aβ production, we first examined its effect on the interaction between APP and BACE1, required for the rate-limiting cleavage step of Aβ production. As a readout of APP-BACE1 interaction, we utilized a previously-published bimolecular fluorescence complementation (BiFC) assay in which APP is tagged with an N-terminal fragment of Venus fluorescent protein (APP:VN), and BACE1 with the complementary C-terminal fragment (BACE:VC)^36^(Fig. 2A). We used flow cytometry to analyze the mean Venus fluorescence intensity in mouse neuroblastoma (N2a) cells co-expressing APP:VN, BACE:VC, and HA-tagged Rab GTPases (for Rab gain-of-function), with Alexa Fluor 647-labeled HA antibody identifying the Rab-expressing cells in which Venus fluorescence was measured (Fig. 2B).

**Figure 2.**
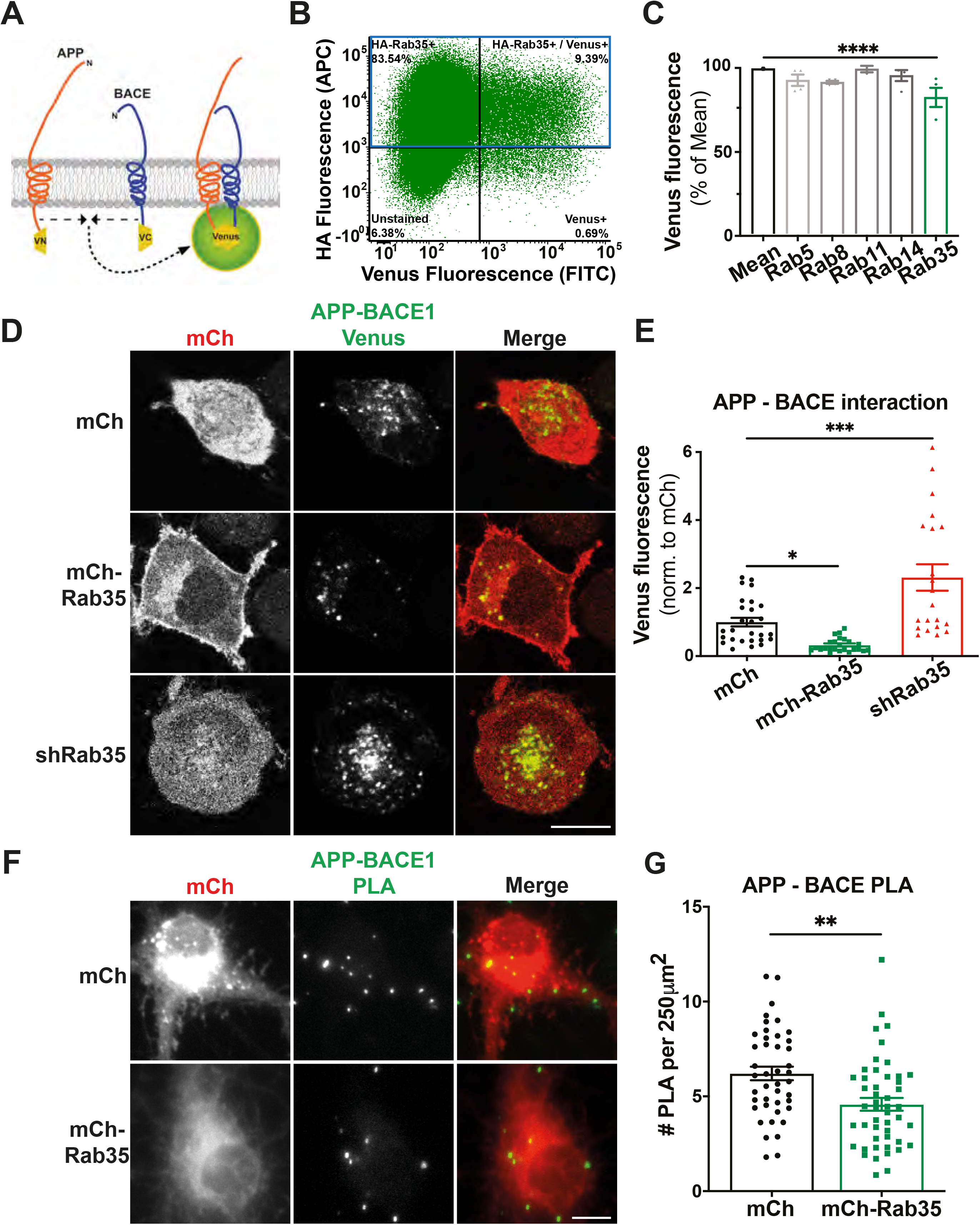
Rab35 is a negative regulator of APP-BACE1 interaction. A) Schematic diagram of bimolecular fluorescence complementation assay, in which APP and BACE1 are tagged with complementary fragments of Venus fluorescent protein (VN and VC, respectively). Reconstitution of Venus fluorescence occurs upon the interaction of APP and BACE1. B) Representative fluorescence-activated cell sorting (FACS) plot of N2a cells co-expressing HA-Rab35 (HA-Rab35+), APP:VN, and BACE:VC (Venus+), as used for analysis shown in panel C. C) Quantification of Venus fluorescence intensity in N2a cells overexpressing the indicated Rab GTPases. Venus fluorescence for each Rab is normalized to Venus fluorescence of the entire Rab-expressing cell population (Mean). Rab35 is a negative regulator of APP-BACE1 interaction (****p< 0.0001; one-way ANOVA, Dunnet post-hoc analysis, n=2 experiments with ~80,000 cells/condition). D-E) Representative images and quantification of Venus fluorescence in N2a cells expressing APP:VN, BACE:VC, and either mCh, mCh-Rab35, or shRab35. Rab35 overexpression and knockdown decrease or increase, respectively, Venus fluorescence when compared to mCh control (*P=0.042, ***P=0.0001; one-way ANOVA, Dunnet post-hoc analysis, n=21-27 cells/condition). F-G) Images and quantification for proximity ligation assay (PLA) to detect endogenous APP-BACE1 interaction in hippocampal neurons expressing mCh or mCh-Rab35. Overexpression of Rab35 decreases PLA puncta density in neuronal cell bodies compared to mCh control. Scale bars: 10 μm. All numeric data represent mean ± SEM (**P =0.001; unpaired student’s t-test, n=43-44 cells/condition).

Compared to average Venus fluorescence across the 16 Rab GTPases investigated (five Rabs associated with endocytic trafficking are shown in Fig. 2C; all Rabs in Fig. S1A), we found that Rab35 gain-of-function decreased the Venus signal by 25% indicating reduced interaction between APP and BACE1 under Rab35 overexpression (Fig. 2C). To confirm these findings, we performed quantitative fluorescence microscopy of N2a cells cotransfected with APP:VN and BACE:VC together with mCh, mCh-Rab35, or mCh coexpressed with an shRNA previously shown to knockdown Rab35 (shRab35) ^37^. Here, Rab35 overexpression again decreased Venus fluorescence by 70% compared to mCh control, suggesting reduced APP-BACE1 interaction, while Rab35 knockdown increased Venus fluorescence by 200% (Fig. 2D-E). Finally, we assessed the impact of Rab35 on endogenous APP-BACE1 interaction in primary rat hippocampal neurons, using the proximity ligation assay (PLA) in neurons transduced either with mCh or mCh-Rab35. Consistent with the BiFC experiments in N2a cells, Rab35 overexpression in primary hippocampal neurons significantly decreased the number of PLA puncta in cell bodies (from 6 to 4.5 puncta/250 μm^2^), representing colocalized APP and BACE1 (Fig. 2F-G). These findings indicate that Rab35 is an important regulator of APP-BACE1 interaction in neurons.

Given that APP is cleaved by BACE1 primarily in endosomes ^19^, our findings suggest that Rab35 may impact APP and/or BACE1 trafficking and localization in the endosomal network. We therefore examined the effects of Rab35 overexpression and knockdown on endogenous APP and BACE1 colocalization with Rab11-positive recycling endosomes in hippocampal neurons. We found that Rab35 overexpression decreased both APP and BACE1 colocalization with Rab11 in the somatodendritic compartment by nearly one-third, while Rab35 knockdown had the opposite effect (Fig. S1A-D). Although Rab35 overexpression decreased the density of Rab11 endosomes (Fig. S1E), its gain- and loss-of-function did not affect endosome size (Fig. S1F), indicating that the observed changes in colocalization were not due to gross effects of Rab35 on the morphology of Rab11 endosomes. Rather, our findings suggest that Rab35 promotes APP and BACE1 sorting out of recycling endosomes.

To understand whether regulation of APP-BACE1 interaction by Rab35 has a functional role in APP processing, we measured levels of APP cleavage products in N2a cells co-transfected with human APP-GFP and either mCherry alone, mCh-Rab35, or mCh + siRNAs to knock down Rab35 (siRab35; see Fig. S2A). Here, overexpression of Rab35 significantly decreased (by ~70%) the levels of APP C-terminal fragments (CTFs) relative to total APP, while knockdown of Rab35 significantly increased CTFs (by ~300%; Fig. 3A-B). A decrease in CTFs was also observed in lysates from human induced pluripotent stem cell (iPSC)-derived cortical neurons lentivirally transduced with mCh or mCh-Rab35 (Fig. 3C-D). To determine whether levels of APP CTFs correlated to similar changes in Aβ production, we measured Aβ40 and Aβ42 peptide concentration in medium collected from these human-derived neurons. Indeed, Rab35 overexpression decreased the levels of both Aβ40 and Aβ42 by 20% (Fig. 3E-F), consistent with an inhibitory role for Rab35 on APP processing through the amyloidogenic pathway.

**Figure 3.**
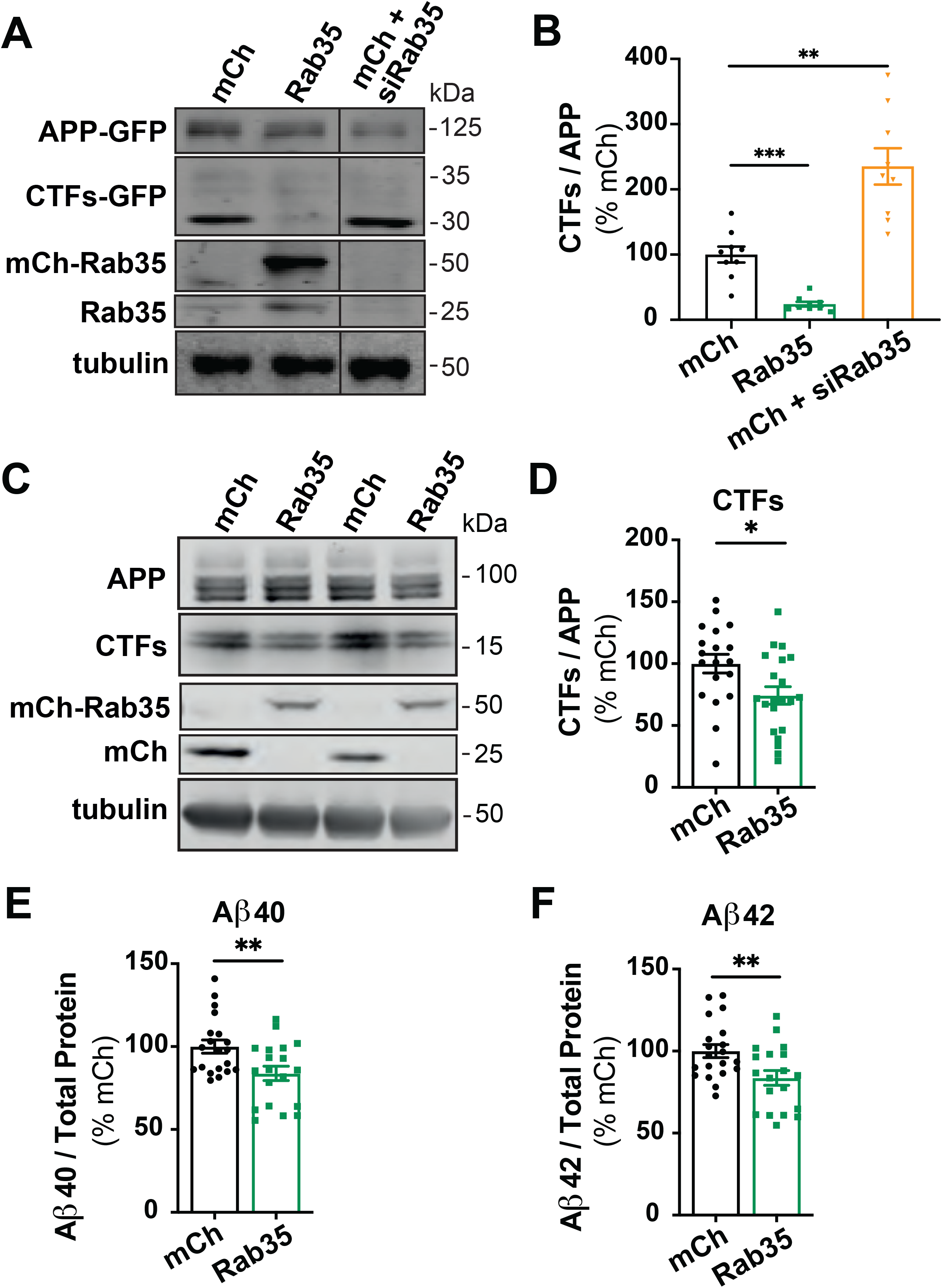
Rab35 expression suppresses amyloidogenic processing of APP. A-C) Representative immunoblots and quantification of APP CTFs from N2a cells expressing mCherry, mCh-Rab35, or mCh + siRNAs against Rab35 (siRab35). Immunoblots were probed for GFP, tubulin, and Rab35, with values normalized to tubulin and expressed as % of mCh control. Overexpression of Rab35 decreases CTFs relative to full-length APP while reduced Rab35 levels exhibits the opposite effects on CTFs (**P_mChCtrl vs mCh-Rab35_=0.0006, **P_mChCtrl vs mCh + siRab35_ =0.0029; one-way ANOVA with Welch’s correction, n=9 samples/condition). C-D) Representative immunoblots and quantification of APP CTFs from iPSC-derived cortical neurons expressing mCh or mCh-Rab35. Blots were probed for APP, tubulin, and mCherry, with values normalized to tubulin and expressed as % of mCh control. Overexpression of Rab35 decreases CTFs relative to full-length APP (*P=0.017; unpaired t-test, n=19-21/condition). E-F) Measurement of Aβ peptides secreted by iPSC-derived human neurons transduced with mCherry or mCh-Rab35. Values are normalized to total protein and expressed as % of mCh control. Rab35 overexpression decreases the levels of both Aβ40 (E) and Aβ42 peptides (F)(**P_Aβ40_=0.008, **P_Aβ42_=0.009; unpaired t-test, n=19-20 samples/ condition). All numeric data represent mean ± SEM.

### Rab35 promotes BACE1 trafficking through the retrograde pathway

Given that Rab35 mediates protein degradation through the endolysosomal pathway ^12,37^, we examined whether Rab35 decreases APP-BACE1 interaction and Aβ production by stimulating the degradation of APP and/or BACE1. Using a previously described cycloheximide (CHX)-chase assay ^37,38^, we found that Rab35 gain- or loss-of-function did not alter the degradation of APP, APP CTFs, or BACE1 in hippocampal neurons (Fig. S2B-F). Since Rab35 also regulates the retrograde trafficking of mannose-6-phosphate receptors from endosomes to the trans-Golgi network (TGN)^13^, we next examined whether Rab35 similarly promotes the retrograde trafficking of APP and/or BACE1. Here, we used an antibody feeding assay in N2a cells, coupled with immunofluorescence microscopy to monitor the colocalization of internalized APP-GFP or FLAG-BACE1 with the TGN marker syntaxin-6 at several timepoints after antibody labeling (see Fig. 4A, S3A). We found that Rab35 overexpression did not alter APP colocalization with syntaxin-6 at any timepoint post-labeling compared to the control condition (Fig. S3B-C). In contrast, Rab35 expression significantly increased the colocalization of internalized BACE1 with syntaxin-6 at later time points (60 and 90 min; Fig. 4B-E), indicating that Rab35 regulates the retrograde trafficking of BACE1, but not APP. To determine whether this trafficking depends on Rab35 activation/GTP binding, we performed the same assay in N2a cells expressing dominant-negative (DN) HA-Rab35. Expression of DN Rab35 either reduced or did not affect BACE1/syntaxin-6 colocalization at the majority of post-labeling timepoints compared to the control condition (Fig. 4B-C). However, DN Rab35 expression was lower than that of WT Rab35 (Fig. S3D), indicating that our results likely underestimate the effects of Rab35 inactivation on BACE1 retrograde trafficking. Nevertheless, these results indicate that Rab35 activation is required for stimulating BACE1 retrograde trafficking.

**Figure 4.**
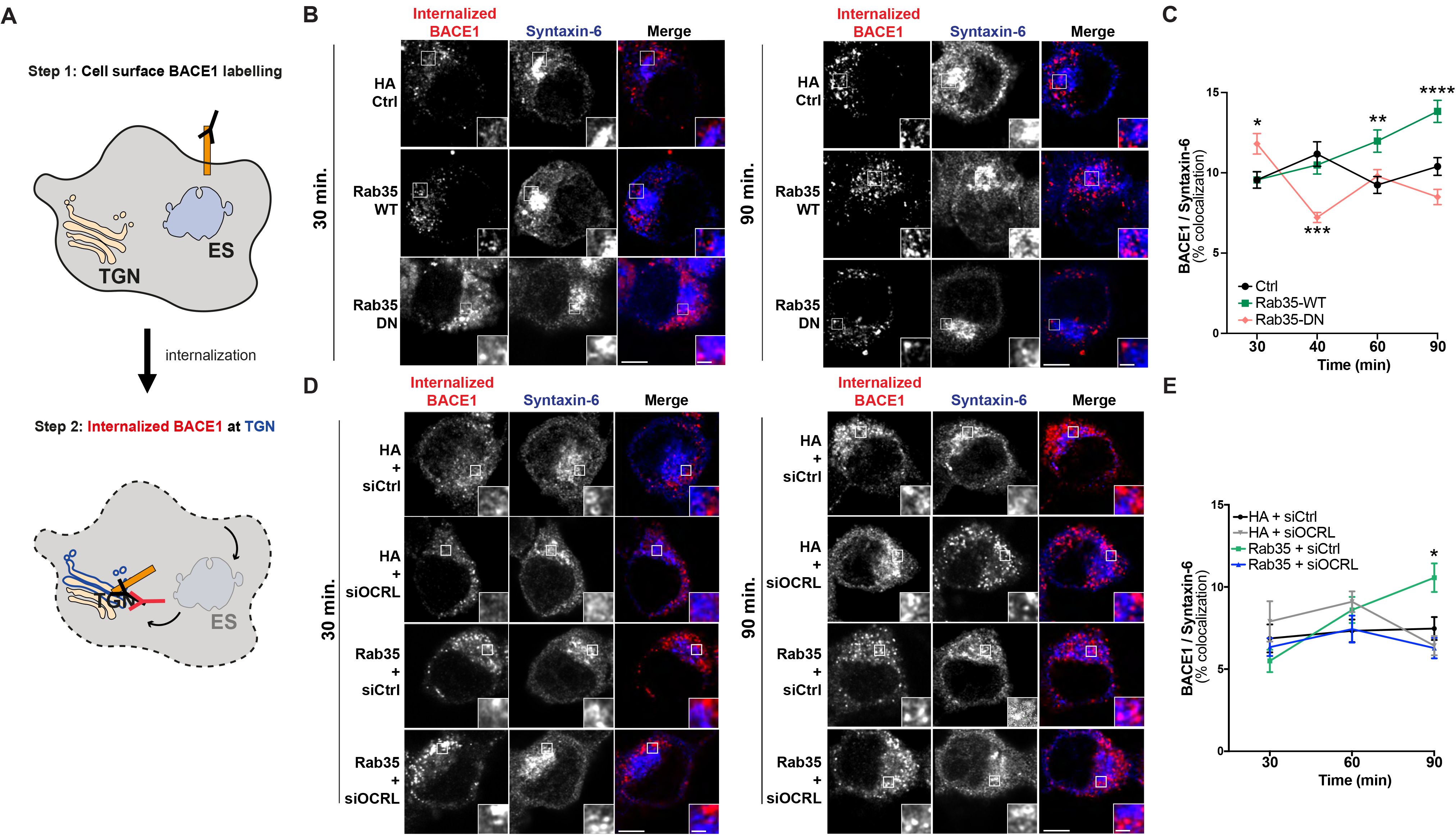
Rab35 stimulates retrograde trafficking of BACE1 via OCRL. A) Schematic representation of BACE1 internalization assay, in which cell-surface BACE1 was labeled with FLAG antibody, cells were incubated for 30, 40, 60, or 90 minutes to allow for BACE1 internalization, and finally, cells were immunostained with syntaxin-6 antibodies to label the TGN. B-C) Representative images and quantification of BACE1 internalization in N2a cells expressing FLAG-BACE1 and either HA Control, HA-Rab35 wild-type (WT), or HA-Rab35 dominant-negative (DN). Internalized BACE1 (red) and syntaxin-6 (blue) are shown at 30 and 90 min time points post-labeling. Compared to vector control, overexpression of WT Rab35 increases colocalization of internalized BACE1 with syntaxin-6 at 60 and 90 min time points, and DN Rab35 prevents this effect (*_PDN-30 min_=0.0392, **P_DN-40 min_=0.0011, **P_WT-60 min_=0.0025, ****P_WT-90 min_ <0.0001; 2-way ANOVA and Sidak post hoc analysis, n=45-155 cells per condition/timepoint. *Time* × *Rab35* interaction F6,1253= 8.002, P<0.0001, overall *Rab35* effect F_2,1253_=10.43, P < 0.0001). D-E) Representative images and quantification of BACE1 internalization in N2a cells expressing FLAG-BACE1 and HA or HA-Rab35 together with control siRNA (siCtrl) or siRNA to knockdown OCRL (siOCRL). Internalized BACE1 (red) and syntaxin-6 (blue) are shown at 30 and 90 min time points post-labeling. Compared to control, overexpression of Rab35 increases colocalization of internalized BACE1 with syntaxin-6 at the 90 min time point, and knockdown of OCRL blocks this effect (*P_HA+siCtrl vs. Rab35+siCtrl_=0.0151, 2-way ANOVA and Tukey’s multiple comparisons test, n=50-61 cells per condition/time point. *Time* × *Condition* interaction F_6,635_=3.861, P=0.009). Scale bars: 5 μm; 1μm for zoomed insets. All numeric data represent mean ± SEM.

The Rab35 effector and lipid phosphatase OCRL (Oculocerebrorenal Syndrome of Lowe Inositol Polyphosphate-5-Phosphatase) is required downstream of Rab35 for the retrograde trafficking of mannose-6-phosphate receptors ^13^. To test whether Rab35-mediated retrograde trafficking of BACE1 also requires OCRL, we performed the same antibody feeding/TGN colocalization assay in the presence of siRNAs against OCRL (siOCRL; see Fig. S3E). While OCRL knockdown alone did not affect BACE1 colocalization with syntaxin-6 compared to control siRNAs, it did prevent the Rab35-mediated increase in BACE1/syntaxin-6 colocalization (Fig. 4D-E). Together, these data indicate that Rab35 stimulates the sorting of BACE1 into the retrograde pathway through its effector OCRL.

To investigate whether Rab35 similarly alters BACE1 trafficking in hippocampal neurons, we measured endogenous APP and BACE1 colocalization with syntaxin-6 following Rab35 overexpression and knockdown. Consistent with our data in N2a cells, APP colocalization with syntaxin-6 was not affected by these manipulations (Fig. S3F-G), while BACE1/syntaxin-6 colocalization was significantly increased by Rab35 overexpression (Fig. S3H-I). This increase was not caused by Rab35-mediated alterations in TGN morphology, as the density and size of syntaxin-6 puncta were unchanged by Rab35 overexpression and knockdown (Fig. S3J-K).

### Rab35 stimulates APP recycling to the plasma membrane

Rab35’s ability to promote the retrograde trafficking of BACE1 to the TGN could be sufficient for reducing APP-BACE1 interaction in the endosomal network. However, Rab35 also facilitates the fast endocytic recycling of proteins (*i.e*. T-cell receptors, β1 integrin) to the plasma membrane (PM) ^14–17^. Stimulating APP and/or BACE1 trafficking into this pathway would also reduce APP-BACE1 interactions in Rab11-positive endosomes. To determine whether Rab35 promotes the fast recycling of APP and/or BACE1, we used another antibody feeding assay to monitor the internalization and PM recycling of APP-GFP and FLAG-BACE1 at four timepoints post-labeling (see Fig. 5A, S4A). Intriguingly, we found that Rab35 overexpression stimulated both APP internalization and recycling to the PM at nearly all postlabeling timepoints (70, 90, and 120 min) compared to the control condition (Fig. 5B-D). Rab35 also stimulated BACE1 internalization at the earlier timepoints post-labeling (60 and 70 min; Fig. S4A-C) but did not alter BACE1 recycling to the PM (Fig. S4B, D). Consistent with these findings, we observed a 40% increase in cell-surface levels of APP, but not BACE1, in N2a cells expressing Rab35 compared to the control condition (Fig. S4E-H).

**Figure 5.**
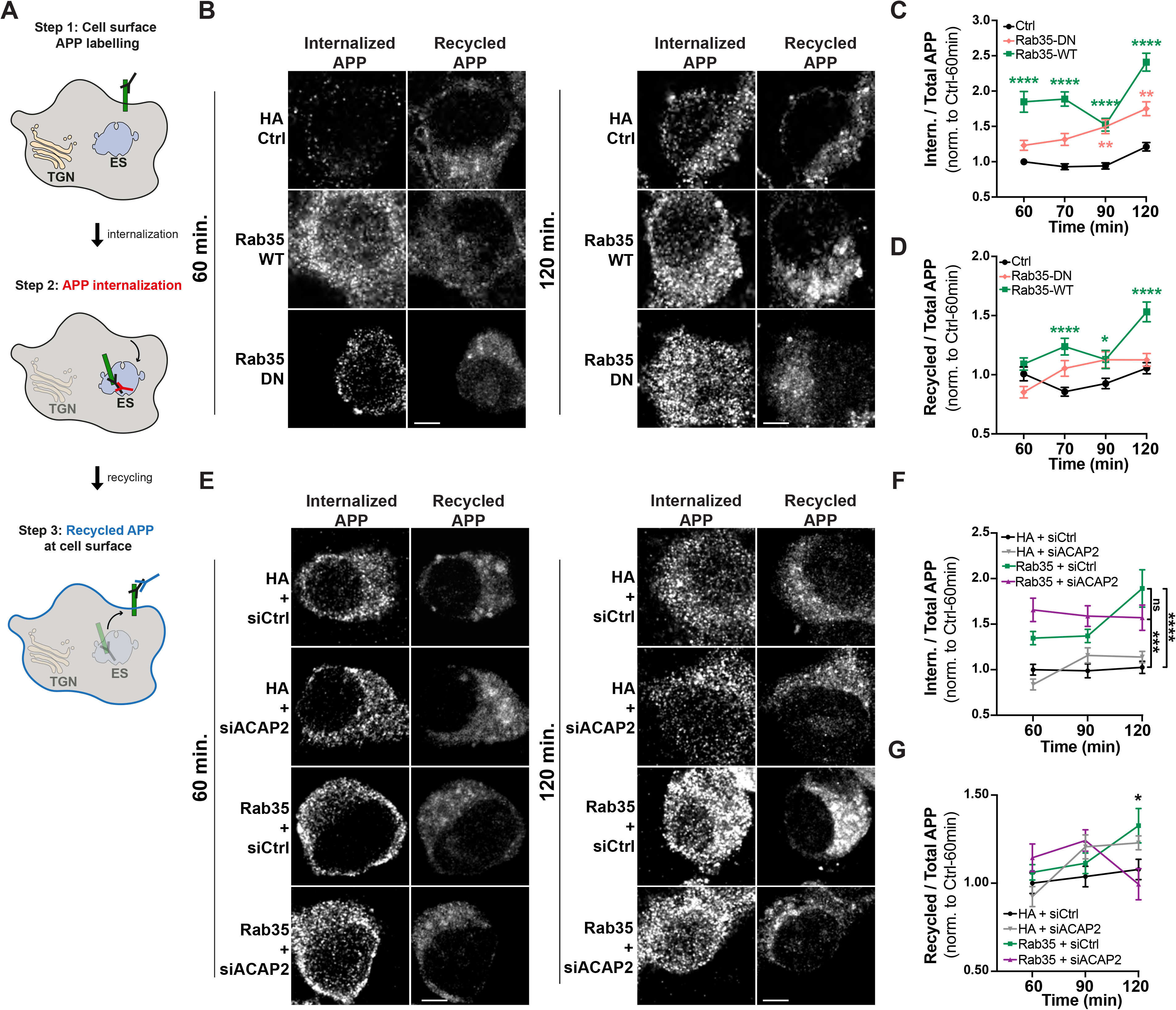
Rab35 stimulates APP recycling to the plasma membrane via ACAP2. A) Schematic representation of the APP recycling assay, in which APP internalization and recycling were assessed by labeling cell-surface APP with 22C11 antibody followed by cell incubation for 60, 70, 90, or 120 minutes, and fixation and immunostaining with secondary antibodies to detect recycled or internalized APP. B-D) Representative images and quantification of APP internalization and recycling in N2a cells expressing APP-GFP and either HA control, HA-Rab35 WT, or HA-Rab35 DN. Internalized and recycled APP are shown at 60 and 90 min time points post-labeling. Compared to control, Rab35 overexpression increases the ratio of internalized (C) and recycled (D) APP over total APP-GFP at multiple timepoints (*P_WT-90 min_=0.0299, **P_DN-90 min_=0.0027, **P_DN-120 min_=0.0017, ****P <0.0001,2-way ANOVA and Sidak post-hoc analysis, n=53-269 cells per condition/timepoint. For APP internalization: *Time* × *Rab35* interaction F_6,1897_=3.002, P=0.0064, overall *Rab35* effect F_2,1897_=118.9. For APP recycling: *Time* × *Rab35* interaction F_6,1897_=3.443, P=0.0022, overall *Rab35* effect F_2,1897_=26.03). E-G) Representative images and quantification of APP internalization and recycling in N2a cells expressing APP-GFP and either HA or HA-Rab35, together with control siRNA (siCtrl) or siRNA to knockdown ACAP2 (siACAP2). Internalized and recycled APP are shown at 60 and 90 min time points post-labeling. F) Compared to control, overexpression of Rab35 increases APP internalization, and ACAP2 knockdown does not alter this effect (***P_HA+siCtrl vs. Rab35+siACAP2_=0.0003, ****P_HA+siCtrl vs. Rab35+siCtrl_<0.0001, 2-way ANOVA and Tukey’s multiple comparisons test, n=20-48 cells per condition/timepoint. *Time* × *Rab35* interaction F_4,310_=2.843, P=0.0244, overall *Rab35/ACAP2* effect F_2,310_=39.12, P< 0.0001). G) Compared to control, overexpression of Rab35 increases APP recycling, and ACAP2 knockdown blocks this effect (*P_HA+siCtrl vs. Rab35+siCtrl_=0.0457, P_HA+siCtrl vs. Rab35+siACAP2_=0.66, 2-way ANOVA with Tukey’s multiple comparisons test, n=20-48 cells per condition/timepoint. *Time* × *Condition* interaction F_4,310_=2.877, P=0.023, overall *Condition* effect F_2,310_=3.279, P=0.039). Scale bars: 5 μm. All numeric data represent mean ± SEM.

To determine whether APP internalization and recycling steps were dependent on Rab35 activation, we also performed this assay in the presence of DN Rab35. As anticipated, the expression of DN Rab35 did not stimulate APP recycling relative to the control condition (Fig. 5B, D), indicating the dependence of this trafficking step on Rab35 activation. Surprisingly, the DN construct did stimulate APP internalization at the 90 and 120 min time points to a similar degree as wild-type Rab35 (Fig. 5B-C), suggesting that Rab35 mediates APP internalization independently of its activation state.

Fast endocytic recycling has been shown to occur through two distinct pathways, mediated by Rab35 effectors OCRL and ACAP2 ^39^. We first tested whether Rab35-mediated APP recycling to the PM relies on OCRL, using the internalization/recycling assay in N2a cells transfected with siRNAs against OCRL (Fig. S5A-C). Interestingly, we found that while OCRL knockdown did not alter Rab35-induced APP internalization, it further increased Rab35-induced APP recycling at the 120 min time point (by ~50%; Fig. S5A, C). These data suggest that Rab35-mediated APP recycling does not occur through OCRL, but that OCRL knockdown frees active Rab35 to interact with other effectors, including the effector responsible for APP recycling, thereby enhancing this trafficking event. We next tested whether this effector is ACAP2, again using our antibody feeding assay in the presence of siRNAs against ACAP2 (Fig. S5D). Here, we found that knockdown of ACAP2 did not alter APP internalization compared to siRNA control, nor lessen the effect of Rab35 on APP internalization (Fig. 5E-F), as expected if this sorting step is a GTP-independent function of Rab35. However, ACAP2 knockdown completely abolished Rab35-enhanced APP recycling to the PM at the 120 min time point (Fig. 5E, G). These data demonstrate that Rab35 stimulates the sorting of APP into the fast recycling pathway, via GTP-independent stimulation of APP internalization and active Rab35/ACAP2-dependent stimulation of APP recycling to the PM. Together, these actions promote APP sorting out of endosomes and increase its accumulation at the PM, thereby decreasing its amyloidogenic processing.

Finally, we tested whether ACAP2, like OCRL, has a role in Rab35-mediated BACE1 retrograde trafficking. Using our antibody feeding/syntaxin-6 colocalization assay to monitor BACE1 retrograde trafficking, we found that knockdown of ACAP2 did not alter the Rab35-mediated increase in BACE1/syntaxin-6 colocalization (Fig. S5E-F). These findings reveal that Rab35 regulates APP and BACE1 trafficking via distinct mechanisms, stimulating the retrograde trafficking of BACE1 to the TGN through OCRL, and the fast recycling of APP to the PM through ACAP2.

### Rab35 counteracts GC-induced amyloidogenic trafficking of APP and BACE1

Although exposure to chronic stress and high GC levels are known to stimulate Aβ production both *in vitro* and *in vivo* ^6,40–42^, their effects on APP and BACE1 intracellular trafficking are largely unexplored. We therefore examined whether high GC levels alter the interaction between APP and BACE1 within the endosomal network, again using the Venus BiFC assay in N2a cells (see Fig. 2). Here, we found that treatment with the synthetic GC dexamethasone (10 μM) significantly increased Venus intensity, indicating that exposure to high GCs promotes APP-BACE1 interaction (Fig. 6A-B). We next examined whether overexpression of Rab35 could block this effect, as predicted if GC-induced downregulation of Rab35 underlies the increased APP-BACE1 interaction. Indeed, Rab35 overexpression blocked the GC-induced increase in Venus intensity (Fig. 6A-B), suggesting that Rab35 prevents this pro-amyloidogenic interaction between APP and BACE1.

**Figure 6.**
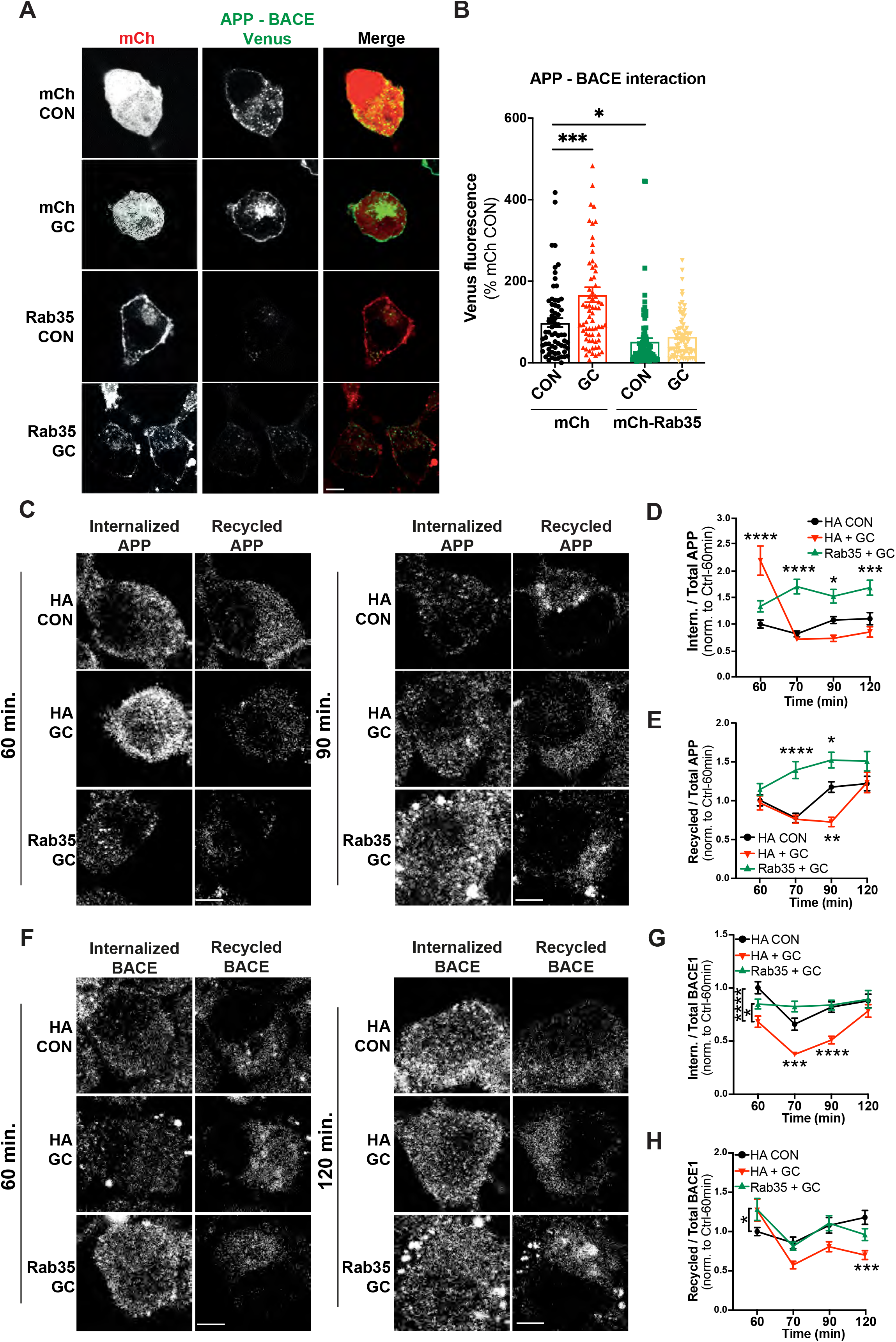
GC-induced amyloidogenic APP/BACE1 trafficking is blocked by Rab35 overexpression. A-B) Representative images and quantification of Venus fluorescence intensity in N2a cells expressing APP:VN, BACE:VC, and either mCh or mCh-Rab35, treated with GC or vehicle control. GC treatment increases Venus intensity in control, but not Rab35-expressing, cells; the latter exhibit an overall decrease in Venus fluorescence (***P_mCh CON vs mCh GC_=0.0004, *P_mCh CON vs mCh-Rab35 CON_=0.028, two-way ANOVA, Tukey post-hoc analysis, n=3 experiments. *Rab35* x *GC* interaction F_1,291_=5.989, P=0.015, overall *Rab35* effect F_1,291_=41.35, P<0.0001, overall *GC* effect F_1,291_=12.19, P=0.0006). Scale bar: 10 μm. C-E) Representative images and quantification of APP internalization and recycling in N2a cells expressing APP-GFP and either HA or HA-Rab35, treated with GC or vehicle control. GC alter APP internalization (D) and recycling (E) kinetics, while Rab35 overexpression prevents these effects (for **D**: *P_HA+DMSO vs HA-Rab35+GC_=0.0119, ***P_HA+DMSO vs HA-Rab35+GC_=0.0009, ****P<0.0001; 2-way ANOVA with Tukey’s post hoc analysis, n=47-77 cells per condition/timepoint. For APP internalization: *Time* × *Condition* interaction F6,780=15.81, P<0.0001. For **E**: ****P_HA+DMSO vs HA-Rab35+GC_<0.0001, *P_HA+DMSO vs HA-Rab35+GC_=0.0136, **P_HA+dmso vs HA+GC_= 0.0011; two-way ANOVA with Tukey’s post hoc analysis, n=47-77 cells per condition/timepoint. For APP recycling: *Time* × *Condition* interaction F_6,759_=3.556, P=0.0013). F-H) Representative images and quantification of BACE1 internalization and recycling in N2a cells expressing FLAG-BACE1 and either HA or HA-Rab35, treated with GCs or vehicle control. GC treatment decreases BACE1 internalization (G) and recycling (H), and these effects are blocked by Rab35 overexpression (for **G**: *P_HA+GC vs HA-Rab35+GC_= 0.0491, ***P_HA+DMSO vs HA-+GC_=0.0003, ****P<0.0001; two-way ANOVA with Tukey’s multiple comparisons test, n=54-76 cells per condition/timepoint. For BACE1 internalization: *Time* × *Condition* interaction F_6,765_=3.385, P=0.0027. For **H**: *P_HA+DMSO vs HA-Rab35+GC_=0.0487, ***P_HA+DMSO vs HA+GC_=0.0009; two-way ANOVA with Tukey’s multiple comparisons test, n=54-76 cells per condition/timepoint. For BACE1 recycling: *Time* × *Condition* interaction F_6,762_=3.504, P=0.0020). Scale bars: 5 μm. All numeric data represent mean ± SEM.

We next tested whether exposure to high GC levels impacts the APP and BACE1 trafficking pathways mediated by Rab35. Using the aforementioned antibody feeding/syntaxin-6 colocalization assay, we found that 24h treatment with GC did not alter BACE1 retrograde trafficking compared to vehicle control (Fig. S6A-B). However, GC treatment significantly altered the kinetics of APP and BACE1 internalization and recycling from the PM, as revealed by the antibody recycling assay. In particular, GC treatment stimulated APP internalization at the 60 min timepoint post-labeling and decreased its recycling back to the PM at the 90 min time point compared to vehicle control (Fig. 6C-E). Additionally, GC decreased BACE1 internalization at multiple time points, as well as BACE1 recycling to the PM (Fig. 6F-H). Remarkably, Rab35 overexpression prevented these GC-induced trafficking deficits and furthermore enhanced APP recycling compared to the control condition (Fig. 6C, E), similar to its actions in the absence of GC (see Fig. 5). Together, these findings demonstrate that GC disrupts the endocytic trafficking of APP and BACE1, resulting in their increased colocalization in endosomes, and that Rab35 overexpression can rescue these GC-driven effects.

## Discussion

Although lifetime stress and exposure to stressful conditions are suggested risk factors for AD, it has been unclear how chronic stress and high GC levels alter cellular trafficking pathways to precipitate Tau and amyloid pathology. Our studies indicate that GC-induced downregulation of the GTPase Rab35, a master regulator of endosomal trafficking, may contribute to both types of pathology. We previously found that Rab35 mediates Tau degradation via the endolysosomal pathway, and that its GC-driven transcriptional suppression leads to Tau accumulation in the hippocampus, inducing synaptic loss and dendritic atrophy ^12^. In the current study, we demonstrate for the first time that Rab35 regulates APP and BACE1 trafficking (Fig. 7), and that its suppression by GCs leads to the accumulation of APP and BACE1 within the endosomal network, increasing Aβ production. Importantly, Rab35 overexpression protects against these GC-driven effects by promoting APP and BACE1 internalization via a putative GTP-independent mechanism, and APP recycling to the plasma membrane via ACAP2 (see Fig. 7). Overall, our findings suggest that downregulation of Rab35 may be a precipitating factor in stress/GC-induced amyloid pathology.

**Figure 7.**
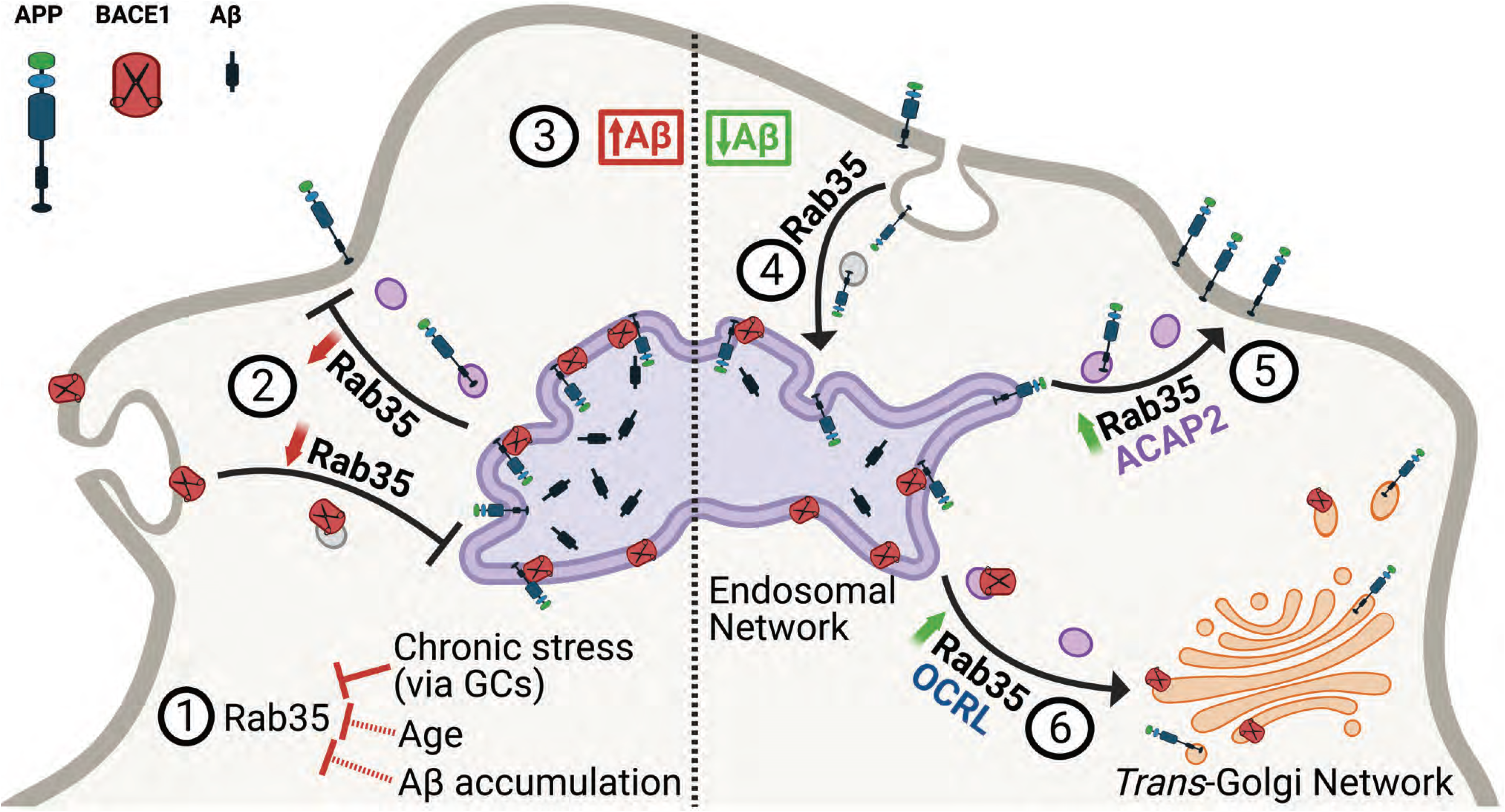
Working model summarizing the interplay between Rab35 and stress/GC on APP trafficking and processing. AD etiopathological factors such as advanced age, Aβ accumulation or exposure to chronic stress and/or high levels of glucocorticoids (GC), lead to a significant reduction of Rab35 levels (1). This reduction inhibits APP and BACE1 recycling with the plasma membrane (2), increasing APP-BACE1 association in the endosomal network and stimulating amyloidogenic APP processing and Aβ production (3). Alternatively, high levels of Rab35 promote APP internalization (4) and recycling to the plasma membrane via the effector ACAP2 (5), and in parallel stimulate BACE1 retrograde trafficking to the trans-Golgi network via the effector OCRL (6). These trafficking events reduce APP-BACE1 interaction within the endosomal network, thereby decreasing production of APP CTFs and Aβ.

APP misprocessing and overproduction of Aβ are widely accepted as the triggering events for AD pathogenesis, underscoring the need to elucidate cellular and molecular mechanisms of amyloidogenic APP processing. Importantly, previous studies show that Aβ is not the only toxic component of the amyloidogenic pathway, as APP β-CTFs (the intracellular product of BACE1 cleavage), have intrinsic neurotoxic properties and can promote Tau hyperphosphorylation and accumulation independently of Aβ, leading to synaptic degeneration and impaired cognition ^43^. Accordingly, and in light of the failures of clinical trials that specifically target Aβ or molecules involved in its cleavage (*e.g*. presenilin)^44^, the focus of more recent studies has shifted to illuminating mechanisms of APP and BACE1 trafficking/interaction in the endosomal network ^45,46^, constituting the rate-limiting step for cleavage of APP into β-CTFs and Aβ peptides. Targeting the molecular modulators of APP and/or BACE1 intracellular trafficking could serve as a promising therapeutic approach against amyloidogenesis ^47^.

Defects in the endolysosomal pathway are the earliest cellular feature of AD ^48,49^, suggesting that dysfunction of endosomal trafficking underlies AD pathogenesis. Indeed, the endosomal network is the major site of Aβ production, and conditions associated with the decreased residence of APP or BACE1 in this subcellular compartment typically inhibit Aβ generation ^20^. Here, we demonstrate that Rab35 decreases the localization of both APP and BACE1 in recycling endosomes (Fig. 7). For APP, this decrease occurs via Rab35-mediated stimulation of APP recycling to the PM through the effector ACAP2, similar to Rab35 stimulation of the fast endocytic recycling of other PM-associated proteins, *e.g*. T-cell receptors and β1 integrin. Not only does this fast recycling pathway decrease APP colocalization with BACE1 in endosomes, but it also boosts levels of APP at the PM (Fig. S4), likely promoting cleavage by α-secretase and thus preventing amyloidogenic processing. For BACE1, the decrease in endosomal sorting appears to result from Rab35-mediated retrograde trafficking to the TGN through the effector OCRL ^13^. Interestingly, two genes identified as risk factors for late-onset AD (LOAD), namely VPS35 and VPS26, encode proteins that regulate the retrograde pathway ^50^. Mutation of these genes is hypothesized to disrupt APP retrograde trafficking to the TGN, increasing the APP-BACE1 interaction in endosomes and therefore stimulating Aβ production ^51,52^. Although we did not see any effect of Rab35 gain-of-function on the retrograde trafficking of APP, we did observe a significant effect on BACE1 retrograde trafficking, thus reducing BACE1 residence time within endosomes as well as its exposure to APP. It is conceivable that Rab35 also regulates BACE1 trafficking through Arf6, another small GTPase previously reported to mediate BACE1 endocytosis from the PM into the endosomal network ^53^. Rab35 and Arf6 have been shown to inhibit one another’s activation in a variety of cellular processes, including cell migration, vesicle secretion, and cytokinesis ^15,54,55^. Thus, Rab35 overexpression could reduce BACE endocytosis via Arf6 inhibition, preventing BACE1 interaction with APP in endosomes. However, we find that cell-surface BACE1 levels are unchanged upon Rab35 overexpression, suggesting that Rab35 does not inhibit BACE1 endocytosis. Future studies should clarify the Arf6-Rab35 signaling relationship in the context of APP and BACE1 trafficking.

An intriguing finding of our study is that Rab35 levels are decreased by several AD etiopathological factors including advanced age, Aβ accumulation, and exposure to chronic stress and/or high levels of the main stress hormones, glucocorticoids (GC). Aging is the greatest risk factor for AD, and studies have linked aging to an increase in amyloidogenic APP processing ^32,33^, although the underlying mechanism(s) of APP misprocessing in aged brain remain unknown. We demonstrate here that Rab35 protein levels are reduced in hippocampus of aged vs. younger rats, and that Rab35 overexpression protects against amyloidogenic APP cleavage in aged animals. Our animal findings are in line with human transcriptomic data in which Rab35 mRNA transcripts are reduced in hippocampus and cortex of older (60-79 year old) individuals compared to younger (20-39 year old) ones. Importantly, reduced Rab35 levels are also observed in hippocampus following Aβ infusion, a non-transgenic experimental model of AD that mimics early AD neuropathology and further stimulates amyloidogenic APP processing ^6,29–31^. In addition to our own observations, other studies have reported that Rab35 levels are reduced under conditions associated with AD risk. For instance, decreased Rab35 levels were observed in the brains of mice expressing the human ApoE4 allele ^56^, the strongest genetic risk factor for LOAD. ApoE4 carriers have a significantly increased probability of developing AD and exhibit greater accumulation of Aβ in their brains compared to carriers of other ApoE alleles^57^. Another recent human study reported decreased Rab35 levels in brain-derived exosomes from athletes following mild traumatic brain injury (TBI)^58^, known to elicit AD-like neuropathology and predispose to AD. Finally, our current *in vitro* and *in vivo* studies show that high GC levels and/or chronic stress reduce Rab35 levels and stimulate the pro-amyloidogenic trafficking of APP and BACE1, while Rab35 overexpression attenuates these effects. These findings are in line with previous studies from us and others demonstrating that chronic stress and high GC levels trigger APP misprocessing and Aβ overproduction ^6,40–42^, and offer a novel, Rab35-linked mechanism to explain how stress/GC precipitate amyloidogenesis. Combined with our previous work showing the critical role of Rab35 in Tau sorting and degradation ^12^, these findings suggest that Rab35 could serve as the molecular link through which chronic stress via GC trigger both major AD pathomechanisms: Aβ overproduction and accumulation of hyperphosphorylated Tau ^6,9,40,59^. Altogether, our findings support multiple roles for Rab35 in the intracellular trafficking of AD-relevant proteins and suggest that downregulation of Rab35 may precipitate amyloidogenesis. These studies highlight Rab35’s potential relevance for AD brain pathology and suggest that additional work investigating its roles in AD-related intracellular pathways could open novel therapeutic avenues for treating AD brain pathology.

## Acknowledgments

This work was supported by NIH grants R01NS080967 and R56AG057560 to C.W., Portuguese Foundation for Science & Technology PhD fellowship PD/BD/105938/2014 to J.V-S., and and PD/BD/135271/2017 to P.G. and the following grants from Foundation for Science and Technology (FCT)-projects UIDB/50026/2020, UIDP/50026/2020 and by the projects NORTE-01-0145-FEDER-000013 and NORTE-01-0145-FEDER-000023, the Project Estratégico co-funded by FCT (PEst-C/SAU/LA0026/2013) and the European Regional Development Fund COMPETE (FCOMP-01-0124-FEDER-037298; POCI-01-0145-FEDER-007038). We would like to thank Dr. Subhojit Roy (University of California, San Diego, USA) for the generous gift of APP:VN and BACE1:VC plasmids, Dr. Andrew Sproul (Stem Cell and Cellular Models Platform, Taub Institute for Research on Alzheimer’s Disease, and the Aging Brain, Columbia University) for providing us with iPSC-derived cortical neurons, and Drs Cristina Mota, Joao Cerqueira and Fernanda Marques (University of Minho) for the hippocampal tissue. We would also like to thank Dr. Ben Wolozin for productive conversations about the manuscript. The human aging RNAseq data were obtained from the Genotype-Tissue Expression (GTEx) Project, supported by the <u>Common Fund</u> of the Office of the Director of the National Institutes of Health, and by NCI, NHGRI, NHLBI, NIDA, NIMH, and NINDS. The data used in this manuscript were obtained from the GTEx Portal on 04/15/20 (dbGaP accession number <u>phs000424.vN.pN</u>).

## Author Contributions

V.Z., J.V-S., I.S., and C.L.W. designed the research; V.Z., J.V-S., M.Z., J.M.S., and P.G. performed experiments and analyzed data; I.S., N. S., and C.L.W. supervised experiments; V.Z., J.V-S., I.S., and C.L.W. wrote the manuscript.

## Materials and Methods

### Primary neurons and cell lines

Primary neuronal cultures were prepared from E18 Sprague Dawley rat embryos and maintained for 14 DIV before use, as described previously ^37^. Neuro2a (N2a) neuroblastoma cells (ATCC CCL-131) were grown in DMEM-GlutaMAX (ThermoFisher) with 10% FBS (Atlanta Biological) and Anti-Anti (ThermoFisher) and kept at 37°C in 5% CO2. During dexamethasone treatment in N2a cells, FBS in the growth media was reduced to 3%. Human iPSC-derived neuronal primary cultures were generated using manual rosette selection and maintained on Matrigel (Corning) ^60^. Concentrated lentiviruses expressing control-sgRNA or hu-APP-sgRNA were made using Lenti-X concentrator (Clontech). The iPSC-derived neuronal cultures were transduced with either control-sgRNA or hu-APP-sgRNA after Accutase splitting and were submitted to puromycin selection the subsequent day. Polyclonal lines were expanded and treated with puromycin for 5 more days before banking. Neuronal differentiation was carried out by plating 165,000 cells/12 well-well in N2/B27 media (DMEM/F12 base) supplemented with BDNF (20 ng/ml) and laminin (1 μg/ml).

### Pharmacological treatments

Pharmacological agents were used in the following concentrations and time courses: cycloheximide (Calbiochem, 0.2 μg/μl, 2, 4 or 8h), dexamethasone (Invivogen, 10 μM, 24 h).

### Lentivirus production, transduction, and DNA transfection

DNA constructs were described previously ^12,37^. APP:VN and BACE:VC constructs were a gift from Dr. Subhojit Roy (University of California, San Diego, USA). Briefly, Rab GTPases were subcloned into pKH3 vector at the EcoRI site to create HA-tagged Rabs. Lentivirus was produced as previously described ^37^. Neurons were transduced with 50–150 μl of lentiviral supernatant per well (12-well plates) or 10–40 μl per coverslip (24-well plates) either at 3 DIV for shRNA transduction or 10 DIV in gain-of-function experiments. Respective controls were transduced on the same day for all experimental conditions. Primary neuronal cultures were collected for immunoblotting or immunocytochemistry at 14 DIV. N2a cells were transfected using Lipofectamine 3000 approximately 24h after plating, according to the manufacturer’s instructions. For the bimolecular fluorescence complementation assay, double transfection of APP:VN and BACE1:VC constructs were performed 48h after transfection with Rab35, to allow for longer expression of the construct. Cells were then fixed and analyzed 18h after the APP:VN and BACE1:VC transfection.

### Flow cytometry

N2a cells were detached using TrypLE Express (Life Technologies) for 5 min at 37°C, resuspended in culture medium, and centrifuged (3,000 rpm, 5 minutes, 4°C). The pellet was washed once with 0.2 mM EDTA and 0.02% BSA in 1x PBS (Flow buffer), centrifuged again, resuspended in 100 μl of flow buffer, then fixed with 4% paraformaldehyde solution for 15 min. Cells were washed (1x PBS) and resuspended in 1.5 % FBS and 0.05% saponin in 1x PBS (permeabilization solution), then placed on a shaker for 30 min. Following centrifugation, the supernatant was discarded, and the cell pellet resuspended in permeabilization solution with anti-HA-tag Alexa(R)-647 (Cell Signaling Technologies) for immunostaining, then placed on a shaker for 90 minutes at 4°C. After washing twice with flow buffer, cells were resuspended in ice-cold Flow Buffer (0,2% FBS, 0.5mM EDTA in PBS), strained through a 35μm nylon mesh to promote single-cell suspensions, and kept on ice. Cells and fluorescence were analyzed by BD Fortessa Cell Analyser and BD FACSDiva software (BD Biosciences). Unstained cells were used as a control for background fluorescence. Far-red (APC) and green (FITC) fluorescence were analyzed, as they marked the HA-tag and Venus fluorescence (APP/BACE1 interaction), respectively. 50,000 events were recorded for each sample, with two samples for each condition. Flow Cytometry data were analyzed using FCS Express 6 (DeNovo Software). Median fluorescence intensity of the Venus (APP/BACE) signal was calculated for the HA+ cells only, thus in double-positive cells. The median Venus fluorescence intensity of each sample was compared to the median Venus fluorescence of all samples, thus comparing each condition to the average of the whole population.

### Proximity ligation assay

Proximity ligation assay (PLA) was performed in primary hippocampal neurons and N2a cells according to the manufacturer’s instructions (Duolink, Sigma). Until the PLA probe incubation step, all manipulations were performed as detailed above for the immunocytochemistry procedure. PLA probes were diluted in blocking solution. The primary antibody pairs used were C1/6.1 (anti-APP, Mouse; Biolegend) and anti-BACE1 (Rabbit, Cell Signaling Technology). All protocol steps were performed at 37°C in a humidity chamber, except for the washing steps. Coverslips were then mounted using Duolink In situ Mounting Media with DAPI.

### Aβ Measurements

Human iPSC-derived neuronal cultures were kept for 3 days post-transduction, after which 50% of the media was changed. Then, conditioned media was collected after 72h, centrifuged at 2,000 rcf for 5 min, and stored at −80°C. Aβ42 and Aβ40 levels were measured using V-PLEX Aβ Peptide Panel 1 (4G8) Kit (MesoScaleDiscovery, MSD) following the manufacturer’s protocol and their concentration was presented as percentage of control levels, after normalization to total protein in the conditioned media (measured using ThermoFisher Scientific BCA assay kit).

### Animals

Male Wistar rats (Charles River Laboratories, France) were maintained under standard laboratory environmental conditions (lights on from 8 a.m to 8 p.m, room temperature 22°C, relative humidity 55%, ad libitum access to food and water). All experimental procedures were approved by the local ethical committee of the University of Minho and the national authority for animal experimentation (DGV9457); all experiments were in accordance with the guidelines for the care and handling of laboratory animals, as described in the Directive 2010/63/EU. 12-month-old animals were randomly divided into the below three groups: control, stressed, Ab-infused animals (N = 8-9 per group). For stressed animals, the chronic unpredictable stress paradigm lasted for 4 weeks and consisted of random application of one of the following stressors (one stressor per day): (i) rocking platform, (ii) air dryer, (iii) cold water, and (iv) overcrowding as previously described^6^. Control, non-stressed animals remain for their home cages during the stress period. At the end of the stress paradigm, all animals were implanted with Alzet miniosmotic pumps for i.c.v. delivery of Aβ1-40 (Eurogentec; 25 μg/200 μl, 0.5μl per hour) or saline for 14 days. For the Aβ1-40 or saline infusion, mini-osmotic pumps (Alzet. Osmotic Pumps, DURECT, 2002 model) and cannulae (Alzet Brain Infusion Kit) were implanted in the left lateral ventricle using the following coordinates from Bregma: −0.6mm anteroposterior, −1.4mm mediolateral, −3.5mm dorsoventral according to Paxinos and Watson ^61^. Pump and cannula implantation were done under anesthesia [75 mg/kg ketamine (Imalgene, Merial) and 1mg/kg medatomidine (Dorbene, Cymedica)]. For aged rat study, young (4 month-old) and aged (22-24 month-old) male Wistar rats were used (N=14 per group). For the Rab35 overexpression experiment, another set of male Wistar rats (17 month-old; Charles River Laboratories, Spain) were randomly divided into two groups (N = 4-8 per group) and were bilaterally injected into the dorsal hippocampus with the AAV8-GFP or AAV8-Rab-GFP virus [coordinates from bregma, according to Paxinos and Watson 50: −3.0 mm anteroposterior (AP), ±1.6 mm mediolateral (ML), and −3.3 mm dorsoventral (DV)] under anesthesia with 75 mg/ kg ketamine (Imalgene, Merial) and 0.5 mg/kg medetomidine (Dorbene, Cymedica) as previously described^12^.

### Western blotting

For western blotting experiments, human iPSC-derived neuronal cultures and N2a cells were collected in Lysis Buffer (50 mm Tris-Base, 150 mm NaCl, 1% Triton X-100, 0.5% deoxycholic acid) with protease inhibitor (Roche) and phosphatase inhibitor cocktails II and III (Sigma) and clarified by centrifugation at high speed (10 min, 20,000 g). Rat hippocampi were homogenized in lysis buffer (50 mm Tris-Base, 150 mm NaCl, 1% Triton X-100, 0.5% deoxycholic acid, 10 mM MgCl_2_) or RIPA buffer with protease inhibitor (Roche) and phosphatase inhibitor cocktails II and III (Sigma) and clarified by centrifugation at high speed (10 min, 20,000 g). Protein concentration was determined using the BCA protein assay kit (ThermoFisher Scientific) and the same amount of protein was used for each condition, which was diluted and denatured in 2× SDS sample buffer (Bio-Rad). Samples were subject to SDS-PAGE, transferred to nitrocellulose membranes using wet (Mini Trans-Blot Cell, Bio-Rad), and probed with the primary antibody in 5% BSA/PBS + 0.1% Tween-20, followed by DyLight 680 or 800 anti-rabbit, anti-mouse (Thermo Scientific) or by HRP-conjugated secondaries (Bio-Rad). Primary antibodies used for western blotting are included in Table 1. Membranes were imaged using an Odyssey Infrared Imager (model 9120, LI-COR Biosciences), and protein intensity was measured using the Image Studio Lite software (LI-COR Biosciences).

**Table 1.**
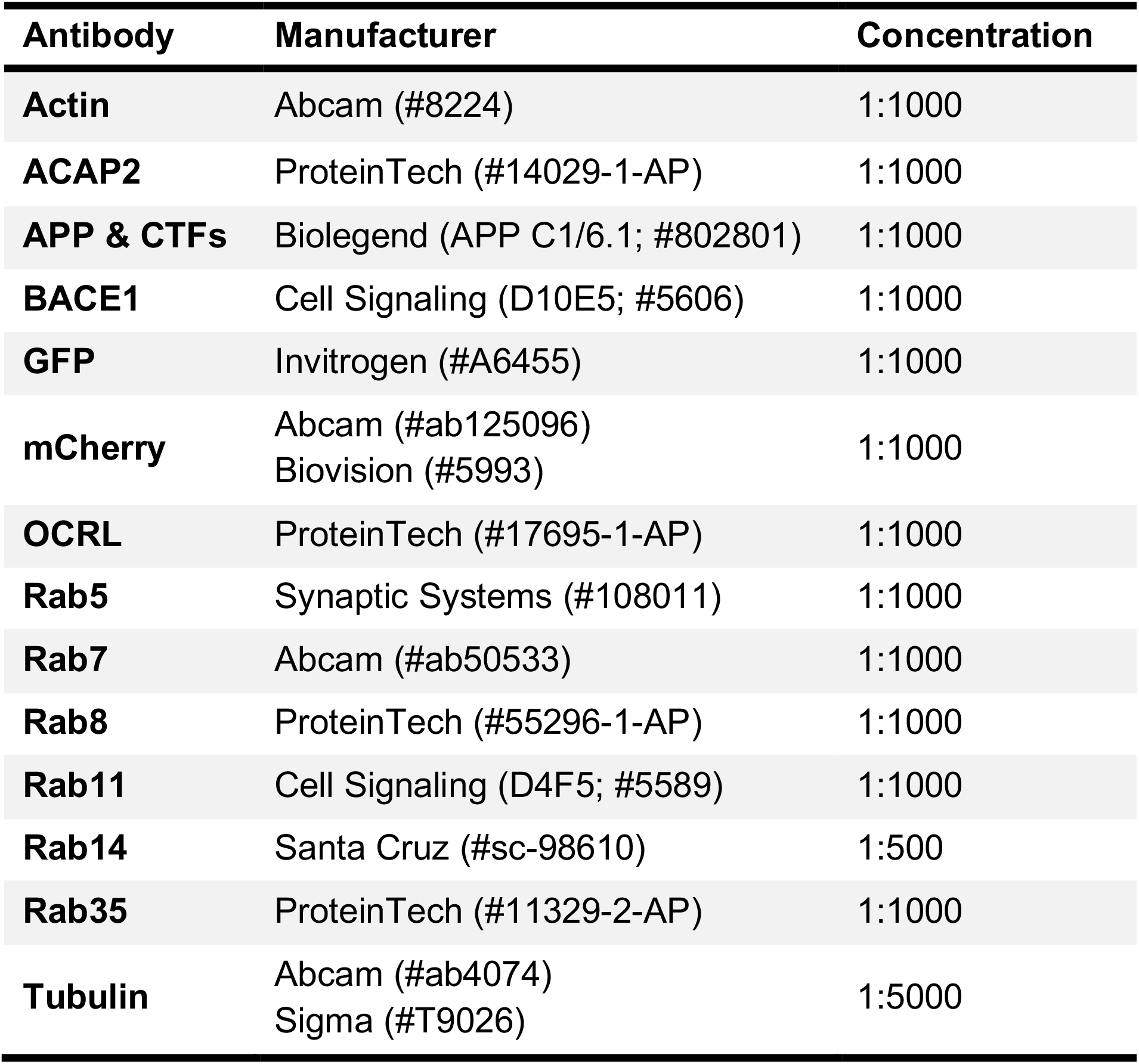
List of antibodies used in this study.

### Immunofluorescence microscopy

Immunofluorescence staining in neurons and N2a cells was performed as previously described ^37^. Briefly, cells were fixed with Lorene’s Fix (60 mM PIPES, 25 mM HEPES, 10 mM EGTA, 2 mM MgCl_2_, 0.12 M sucrose, 4% formaldehyde) for 15 min, and primary and secondary antibody incubations were performed in blocking buffer (2% glycine, 2% BSA, 0.2% gelatin, 50 mM NH4Cl in 1× PBS) for 1 h at room temperature. Images were acquired using a Zeiss LSM 800 confocal microscope equipped with Airyscan module, using a 63× objective (Plan-Apochromat, NA 1.4; for neurons and N2a cell imaging).

### Neuronal image analyses

Images were analyzed and processed using the Fiji software. PLA puncta were counted using the Multi-point tool, and cell area was measured with Polygon selection tool. Colocalization analysis between APP (22C11, Millipore) or BACE1 (Cell Signaling) and intracellular compartments (Rab11, Cell Signaling; Syntaxin-6, Synaptic Systems) in neurons was determined using the JACoP plugin, in order to obtain the Mander’s coefficient corresponding to the fraction of APP or BACE1 colocalized with each compartment.

### Retrograde trafficking assay

N2a cells were co-transfected with APP-GFP or FLAG-BACE1 and HA or HA-Rab35 constructs. Approximately 48 hours after transfection, cells were starved in serum-free DMEM for 30 minutes. Cells were then incubated for 30 min at 4°C with 22C11 antibody (anti-N-terminus of APP, Millipore) or anti-FLAG (Millipore). Antibodies were diluted 1:100 in complete medium + 1M HEPES. Following antibody incubation, cells were washed with complete medium + HEPES and either immediately fixed with Lorene’s fixative for 15 min, or incubated at 37°C for 10, 30, or 60 min and then fixed, followed by washing with 1X PBS. For immunostaining, cells were permeabilized using Triton X-100 and coverslips were immunostained with the following primary + secondary antibody pairs: internalized anti-22C11 or anti-FLAG + goat-anti-mouse Alexa Fluor-568; anti-Syntaxin-6 (Synaptic Systems) + Alexa Fluor-647 to tag the trans-Golgi network (TGN); and anti-BACE1 (for BACE1-transfected conditions only, Cell Signaling) + Alexa Fluor-488 to tag total BACE1. Cells overexpressing Rab35 were detected using an anti-HA primary antibody (data not shown; Rabbit, Cell Signaling; Mouse, Biolegend) + Alexa Fluor-405 secondary antibody. For analysis, Fiji/ImageJ was used to outline each transfected cell and clear the background. Colocalization analysis between APP or BACE1 and the TGN was determined using the JACoP plugin, and the Mander’s coefficient was used for reporting the fraction of APP or BACE1 colocalized with the TGN.

### Recycling assay

N2a cells were co-transfected with APP-GFP or FLAG-BACE1 and HA or HA-Rab35 constructs. Approximately 48 hours after transfection, cells were starved in serum-free DMEM for 30 minutes. Cells were then incubated for 30 min at 4°C with 22C11 antibody (anti-N-terminus of APP, Millipore) or anti-FLAG (Millipore). Following antibody pulse, coverslips were washed with complete medium + HEPES and incubated with goat-anti-mouse unconjugated antibody (1:50; Invitrogen) for 30 min at 4°C to allow for APP or BACE internalization. Antibodies were diluted in complete medium + 1M HEPES. Coverslips were washed with complete medium + HEPES and fixed with Lorene’s fixative for 15 min or incubated at 37°C for 10, 30, or 60 min and then fixed, followed by washing with 1X PBS. Coverslips were immunostained with goat-anti-mouse Alexa Fluor-568 for 1hr at RT prior to cell permeabilization to mark recycled 22C11 or FLAG antibodies, and any remaining surface antibody was blocked using goat-anti-mouse unconjugated antibody (1:50, 30 min at RT). Following cell permeabilization, internalized 22C11 or FLAG was marked using goat-anti-mouse Alexa Fluor-647 secondary antibody. Total BACE1 and HA were tagged as in the retrograde trafficking assay, and antibody concentrations are the same as above. Internalization and recycling of APP and BACE1 were determined using Fiji/ImageJ by outlining each cell and normalizing the fluorescence of (1) internalized APP or BACE1 and (2) recycled APP or BACE1 to total APP or BACE1. These values were then normalized to the first control timepoint for APP or BACE1 to determine change over time.

### Steady-state surface protein measurement

N2a cells were co-transfected as in the retrograde trafficking and recycling assays. Approximately 48 hours after transfection, surface APP and BACE1 were labeled with anti-22C11 or anti-FLAG antibodies prior to cell permeabilization. Following permeabilization, coverslips were immunostained for HA (APP-GFP transfected cells) or HA and BACE1 as described for the retrograde trafficking assay. Fiji/ImageJ was used to determine surface APP and BACE1 by outlining each transfected cell and normalizing the fluorescence of surface APP or BACE1 to total APP or BACE1.

### Bioinformatics analysis

RNA sequencing data across age groups were collected and produced by the Genotype-Tissue Expression (GTEx) project (https://gtexportal.org/home/). Rab35 transcripts per million were normalized to IPO8 transcripts per million for each sample using Microsoft Excel. IPO8 was chosen as a reference gene because it was the most stable gene tested from the GTEx dataset, using RefFinder ^34^.

### Statistical analysis

Graphing and statistics analysis was performed using Prism (GraphPad). Shapiro–Wilk normality test was used to determine whether data sets were modeled by a normal distribution. Unpaired, two-tailed t-tests, one-way ANOVA, or two-way ANOVAs were used with values of P < 0.05 being considered as significantly different.

## References

1 Querfurth, H. W. & LaFerla, F. M. Alzheimer’s disease. N Engl J Med 362, 329–344, doi:10.1056/NEJMra0909142 (2010).

2 Mah, L., Szabuniewicz, C. & Fiocco, A. J. Can anxiety damage the brain? Current opinion in psychiatry 29, 56–63, doi:10.1097/YCO.0000000000000223 (2016).

3 Machado, A. et al. Chronic stress as a risk factor for Alzheimer’s disease. Rev Neurosci 25, 785–804, doi:10.1515/revneuro-2014-0035 (2014).

4 Mravec, B., Horvathova, L. & Padova, A. Brain Under Stress and Alzheimer’s Disease. Cell Mol Neurobiol 38, 73–84, doi:10.1007/s10571-017-0521-1 (2018).

5 Vyas, S. et al. Chronic Stress and Glucocorticoids: From Neuronal Plasticity to Neurodegeneration. Neural plasticity 2016, 6391686, doi:10.1155/2016/6391686 (2016).

6 Catania, C. et al. The amyloidogenic potential and behavioral correlates of stress. Mol Psychiatry 14, 95–105, doi:10.1038/sj.mp.4002101 (2009).

7 Sotiropoulos, I. et al. Stress acts cumulatively to precipitate Alzheimer’s disease-like tau pathology and cognitive deficits. J Neurosci 31, 7840–7847, doi:10.1523/JNEUROSCI.0730-11.2011 (2011).

8 Kulstad, J. J. et al. Effects of chronic glucocorticoid administration on insulin-degrading enzyme and amyloid-beta peptide in the aged macaque. J Neuropathol Exp Neurol 64, 139–146 (2005).

9 Sotiropoulos, I. et al. Glucocorticoids trigger Alzheimer disease-like pathobiochemistry in rat neuronal cells expressing human tau. J Neurochem 107, 385–397, doi:10.1111/j.1471-4159.2008.05613.x (2008).

10 Lopes, S. et al. Tau Deletion Prevents Stress-Induced Dendritic Atrophy in Prefrontal Cortex: Role of Synaptic Mitochondria. Cereb Cortex, doi:10.1093/cercor/bhw057 (2016).

11 Pinheiro, S. et al. Tau Mislocation in Glucocorticoid-Triggered Hippocampal Pathology. Mol Neurobiol, doi:10.1007/s12035-015-9356-2 (2015).

12 Vaz-Silva, J. et al. Endolysosomal degradation of Tau and its role in glucocorticoid-driven hippocampal malfunction. EMBO J, doi:10.15252/embj.201899084 (2018).

13 Cauvin, C. et al. Rab35 GTPase Triggers Switch-like Recruitment of the Lowe Syndrome Lipid Phosphatase OCRL on Newborn Endosomes. Curr Biol 26, 120–128, doi:10.1016/j.cub.2015.11.040 (2016).

14 Argenzio, E. et al. CLIC4 regulates cell adhesion and beta1 integrin trafficking. J Cell Sci 127, 5189–5203, doi:10.1242/jcs.150623 (2014).

15 Allaire, P. D. et al. Interplay between Rab35 and Arf6 controls cargo recycling to coordinate cell adhesion and migration. J Cell Sci 126, 722–731, doi:10.1242/jcs.112375 (2013).

16 Patino-Lopez, G. et al. Rab35 and its GAP EPI64C in T cells regulate receptor recycling and immunological synapse formation. J Biol Chem 283, 18323–18330, doi:10.1074/jbc.M800056200 (2008).

17 Kouranti, I., Sachse, M., Arouche, N., Goud, B. & Echard, A. Rab35 regulates an endocytic recycling pathway essential for the terminal steps of cytokinesis. Curr Biol 16, 1719–1725, doi:10.1016/j.cub.2006.07.020 (2006).

18 Zou, L. et al. Receptor tyrosine kinases positively regulate BACE activity and Amyloidbeta production through enhancing BACE internalization. Cell Res 17, 389–401, doi:10.1038/cr.2007.5 (2007).

19 Haass, C., Kaether, C., Thinakaran, G. & Sisodia, S. Trafficking and proteolytic processing of APP. Cold Spring Harbor perspectives in medicine 2, a006270, doi:10.1101/cshperspect.a006270 (2012).

20 Small, S. A. S.-S., S.; Mayeux, R.; Petsko, G.A. Endosomal traffick jams represent a pathogenic hub and therapeutic target in Alzheimer’s disease. Trends in Neurosciences 40, 592–602 (2017).

21 Ubelmann, F. et al. Bin1 and CD2AP polarise the endocytic generation of beta-amyloid. EMBO Rep 18, 102–122, doi:10.15252/embr.201642738 (2017).

22 Udayar, V. et al. A paired RNAi and RabGAP overexpression screen identifies Rab11 as a regulator of beta-amyloid production. Cell reports 5, 1536–1551, doi:10.1016/j.celrep.2013.12.005 (2013).

23 Ginsberg, S. D. et al. Regional selectivity of rab5 and rab7 protein upregulation in mild cognitive impairment and Alzheimer’s disease. J Alzheimers Dis 22, 631–639, doi:10.3233/JAD-2010-101080 (2010).

24 Goncalves, S. A. & Outeiro, T. F. Traffic jams and the complex role of alpha-Synuclein aggregation in Parkinson disease. Small Gtpases 8, 78–84, doi:10.1080/21541248.2016.1199191 (2017).

25 Masaracchia, C. et al. Membrane binding, internalization, and sorting of alpha-synuclein in the cell. Acta Neuropathol Commun 6, 79, doi:10.1186/s40478-018-0578-1 (2018).

26 Frautschy, S. A., Yang, F., Calderon, L. & Cole, G. M. Rodent models of Alzheimer’s disease: rat A beta infusion approaches to amyloid deposits. Neurobiol Aging 17, 311–321, doi:10.1016/0197-4580(95)02073-x (1996).

27 Geula, C. et al. Aging renders the brain vulnerable to amyloid beta-protein neurotoxicity. Nat Med 4, 827–831, doi:10.1038/nm0798-827 (1998).

28 Stephan, A. & Phillips, A. G. A case for a non-transgenic animal model of Alzheimer’s disease. Genes Brain Behav 4, 157–172, doi:10.1111/j.1601-183X.2004.00113.x (2005).

29 Heredia, L. et al. Deposition of amyloid fibrils promotes cell-surface accumulation of amyloid beta precursor protein. Neurobiol Dis 16, 617–629, doi:10.1016/j.nbd.2004.04.015 (2004).

30 Lorenzo, A. et al. Amyloid beta interacts with the amyloid precursor protein: a potential toxic mechanism in Alzheimer’s disease. Nat Neurosci 3, 460–464, doi:10.1038/74833 (2000).

31 Davis-Salinas, J., Saporito-Irwin, S. M., Cotman, C. W. & Van Nostrand, W. E. Amyloid beta-protein induces its own production in cultured degenerating cerebrovascular smooth muscle cells. J Neurochem 65, 931–934, doi:10.1046/j.1471-4159.1995.65020931.x (1995).

32 Cisternas, P. et al. New Insights into the Spontaneous Human Alzheimer’s Disease-Like Model Octodon degus: Unraveling Amyloid-beta Peptide Aggregation and Age-Related Amyloid Pathology. J Alzheimers Dis 66, 1145–1163, doi:10.3233/JAD-180729 (2018).

33 Kimura, N. et al. Dynein Dysfunction Reproduces Age-Dependent Retromer Deficiency: Concomitant Disruption of Retrograde Trafficking Is Required for Alteration in beta-Amyloid Precursor Protein Metabolism. Am J Pathol 186, 1952–1966, doi:10.1016/j.ajpath.2016.03.006 (2016).

34 Xie, F., Xiao, P., Chen, D., Xu, L. & Zhang, B. miRDeepFinder: a miRNA analysis tool for deep sequencing of plant small RNAs. Plant Mol Biol, doi:10.1007/s11103-012-9885-2 (2012).

35 Sotiropoulos, I., Silva, J. M., Gomes, P., Sousa, N. & Almeida, O. F. X. Stress and the Etiopathogenesis of Alzheimer’s Disease and Depression. Adv Exp Med Biol 1184, 241–257, doi:10.1007/978-981-32-9358-8_20 (2019).

36 Das, U. et al. Visualizing APP and BACE-1 approximation in neurons yields insight into the amyloidogenic pathway. Nat Neurosci 19, 55–64, doi:10.1038/nn.4188 (2016).

37 Sheehan, P., Zhu, M., Beskow, A., Vollmer, C. & Waites, C. L. Activity-Dependent Degradation of Synaptic Vesicle Proteins Requires Rab35 and the ESCRT Pathway. J Neurosci 36, 8668–8686, doi:10.1523/JNEUROSCI.0725-16.2016 (2016).

38 Miranda, A. M. et al. Neuronal lysosomal dysfunction releases exosomes harboring APP C-terminal fragments and unique lipid signatures. Nature communications 9, 291, doi:10.1038/s41467-017-02533-w (2018).

39 Mrozowska, P. S. & Fukuda, M. Regulation of podocalyxin trafficking by Rab small GTPases in 2D and 3D epithelial cell cultures. J Cell Biol 213, 355–369, doi:10.1083/jcb.201512024 (2016).

40 Green, K. N., Billings, L. M., Roozendaal, B., McGaugh, J. L. & LaFerla, F. M. Glucocorticoids increase amyloid-beta and tau pathology in a mouse model of Alzheimer’s disease. J Neurosci 26, 9047–9056, doi:10.1523/JNEUROSCI.2797-06.2006 (2006).

41 Jeong, Y. H. et al. Chronic stress accelerates learning and memory impairments and increases amyloid deposition in APPV717I-CT100 transgenic mice, an Alzheimer’s disease model. FASEB J 20, 729–731, doi:10.1096/fj.05-4265fje (2006).

42 Srivareerat, M., Tran, T. T., Alzoubi, K. H. & Alkadhi, K. A. Chronic psychosocial stress exacerbates impairment of cognition and long-term potentiation in beta-amyloid rat model of Alzheimer’s disease. Biol Psychiatry 65, 918–926, doi:10.1016/j.biopsych.2008.08.021 (2009).

43 Moore, S. et al. APP metabolism regulates tau proteostasis in human cerebral cortex neurons. Cell reports 11, 689–696, doi:10.1016/j.celrep.2015.03.068 (2015).

44 Huang, L. K., Chao, S. P. & Hu, C. J. Clinical trials of new drugs for Alzheimer disease. J Biomed Sci 27, 18, doi:10.1186/s12929-019-0609-7 (2020).

45 Tan, J. Z. A. & Gleeson, P. A. The role of membrane trafficking in the processing of amyloid precursor protein and production of amyloid peptides in Alzheimer’s disease. Biochim Biophys Acta Biomembr 1861, 697–712, doi:10.1016/j.bbamem.2018.11.013 (2019).

46 Bera, S. et al. AP-2 reduces amyloidogenesis by promoting BACE1 trafficking and degradation in neurons. EMBO Rep 21, e47954, doi:10.15252/embr.201947954 (2020).

47 Sun, J. & Roy, S. The physical approximation of APP and BACE-1: A key event in alzheimer’s disease pathogenesis. Dev Neurobiol 78, 340–347, doi:10.1002/dneu.22556 (2018).

48 Nixon, R. A. Endosome function and dysfunction in Alzheimer’s disease and other neurodegenerative diseases. Neurobiol Aging 26, 373–382, doi:S0197-4580(04)00294-5 [pii] 10.1016/j.neurobiolaging.2004.09.018 (2005).

49 Orr, M. E. & Oddo, S. Autophagic/lysosomal dysfunction in Alzheimer’s disease. Alzheimers Res Ther 5, 53, doi:10.1186/alzrt217 (2013).

50 Small, S. A. Retromer sorting: a pathogenic pathway in late-onset Alzheimer disease. Archives of neurology 65, 323–328, doi:10.W01/archneurol.2007.64 (2008).

51 Sullivan, C. P. et al. Retromer disruption promotes amyloidogenic APP processing. Neurobiol Dis 43, 338–345, doi:10.1016/j.nbd.2011.04.002 (2011).

52 Bhalla, A. et al. The location and trafficking routes of the neuronal retromer and its role in amyloid precursor protein transport. Neurobiol Dis 47, 126–134, doi:10.1016/j.nbd.2012.03.030 (2012).

53 Sannerud, R. et al. ADP ribosylation factor 6 (ARF6) controls amyloid precursor protein (APP) processing by mediating the endosomal sorting of BACE1. Proc Natl Acad Sci U S A 108, E559–568, doi:10.1073/pnas.1100745108 (2011).

54 Kobayashi, H. & Fukuda, M. Rab35 regulates Arf6 activity through centaurin-beta2 (ACAP2) during neurite outgrowth. J Cell Sci 125, 2235–2243, doi:10.1242/jcs.098657 (2012).

55 Sheehan, P. & Waites, C. L. Coordination of synaptic vesicle trafficking and turnover by the Rab35 signaling network. Small Gtpases, 1–10, doi:10.1080/21541248.2016.1270392 (2017).

56 Peng, K. Y. et al. Apolipoprotein E4 genotype compromises brain exosome production. Brain 142, 163–175, doi:10.1093/brain/awy289 (2019).

57 Huang, Y. A., Zhou, B., Wernig, M. & Sudhof, T. C. ApoE2, ApoE3, and ApoE4 Differentially Stimulate APP Transcription and Abeta Secretion. Cell 168, 427–441 e421, doi:10.1016/j.cell.2016.12.044 (2017).

58 Goetzl, E. J. et al. Neuron-Derived Exosome Proteins May Contribute to Progression From Repetitive Mild Traumatic Brain Injuries to Chronic Traumatic Encephalopathy. Front Neurosci 13, 452, doi:10.3389/fnins.2019.00452 (2019).

59 Carroll, J. C. et al. Chronic stress exacerbates tau pathology, neurodegeneration, and cognitive performance through a corticotropin-releasing factor receptor-dependent mechanism in a transgenic mouse model of tauopathy. J Neurosci 31, 14436–14449, doi:10.1523/JNEUROSCI.3836-11.2011 (2011).

60 Topol, A., Tran, N. N. & Brennand, K. J. A guide to generating and using hiPSC derived NPCs for the study of neurological diseases. J Vis Exp, e52495, doi:10.3791/52495 (2015).

61 Paxinos, G., Watson, C., Pennisi, M. & Topple, A. Bregma, lambda and the interaural midpoint in stereotaxic surgery with rats of different sex, strain and weight. J Neurosci Methods 13, 139–143, doi:10.1016/0165-0270(85)90026-3 (1985).

